# Analysis of half a billion datapoints across ten machine-learning algorithms identify key determinants of insulin gene transcription

**DOI:** 10.1101/2021.03.27.437353

**Authors:** Wilson K.M. Wong, Vinod Thorat, Mugdha V. Joglekar, Charlotte X. Dong, Hugo Lee, Yi Vee Chew, Adwait Bhave, Wayne J. Hawthorne, Feyza Engin, Aniruddha Pant, Louise T. Dalgaard, Sharda Bapat, Anandwardhan A. Hardikar

## Abstract

Machine learning (ML)-workflows enable unprejudiced/robust evaluation of complex datasets. Here, we analyzed over 490,000,000 data points to compare 10 different ML-workflows in a large (N=11,652) training dataset of human pancreatic single-cell (sc-)transcriptomes to identify genes associated with the presence or absence of insulin transcript(s). Prediction accuracy/sensitivity of each ML-workflow were tested in a separate validation dataset (N=2,913). Ensemble ML-workflows, in particular Random Forest ML-algorithm delivered high predictive power (AUC=0.83) and sensitivity (0.98), compared to other algorithms. The transcripts identified through these analyses also demonstrated significant correlation with insulin in bulk RNA-seq data from human islets. The top-10 features, (including *IAPP, ADCYAP1, LDHA* and *SST*) common to the three Ensemble ML-workflows were significantly dysregulated in scRNA-seq datasets from Ire-1α^β-/-^ mice that demonstrate dedifferentiation of pancreatic β-cells in a model of type 1 diabetes (T1D) and in pancreatic single cells from individuals with type 2 Diabetes (T2D). Our findings provide direct comparison of ML-workflows in big data analyses, identify key determinants of insulin transcription and provide workflows for future analyses.

## Introduction

Recent years have witnessed a surge in single-cell transcriptomic technologies; many already generating newer data and insights to address specific biological questions. Machine learning (ML) algorithms offer an unbiased mathematical workflow that facilitates the identification of complex relationships across variables. ML workflows involve an orderly set of instructions using automated, unbiased ‘learning’ processes usually targeted towards developing (training) a model that can be validated in a separate (test) dataset[1]. One goal of ML algorithms is to analyze big data to identify variables that cannot be recognized through conventional biostatistical techniques, and enhance development of predictive algorithms[2, 3].

Currently, several ML algorithms are available to researchers handling big data in omics-based high content analyses. These can be broadly divided into two categories: supervised and unsupervised algorithms[4]. Supervised methods (such as decision tree) derive relationships between one dependent and multiple independent variables using a training set and then apply that knowledge in the testing set for predictive/efficacy analysis. Unsupervised methods derive patterns/data clusters amongst all available variables. ML algorithms have been used to unravel patterns/clustering in high-density transcriptome analyses[5] or to build associations[6] or for predictions in several biological processes such as determining DNA methylation states in single cells[7], identifying signatures of lipid or metabolite species[8] or microRNAs[9] in predicting transition from gestational diabetes to type 2 diabetes as well as in genetic studies[10]. There are multiple ML algorithms available and it may present a challenge to select the most appropriate method for a particular dataset to answer a specific question. We, therefore, decided to compare different ML methodologies to (i) rank different ML methods for their performance on a large dataset (of 490,855,065 scRNA-sequencing data points) and (ii) understand the most important variables associated with insulin transcription.

Previous studies[11-14] from several laboratories have identified master regulatory transcription factors that regulate the embryonic development of insulin-producing islet β-cells. Although transcription factor-mediated insulin transcription regulation is a well-known mechanism during the development of insulin-producing cells, it is also recognized that active genes localized on different chromosomal regions can dynamically regulate gene transcription in post-natal life[15]. One approach to identify genes associated with insulin gene transcription is through single-cell (sc)RNA-seq-based big data analysis.

Here, we examined the performance of 10 different ML algorithms in a curated human pancreatic single-cell sequencing dataset of 490,855,065 data points (N=14,565 single cells and 33,701 expressed gene features). The aims of this study were (i) to provide a comparative account of the predictive potential of 10 different commonly used ML workflows (**Supplementary Table 1**), and (ii) to use existing scRNA-seq datasets in identifying genes (variables) associated with or important for determining insulin transcript-containing cells.

## Results

### Machine learning (ML) algorithms yield varying performance outputs

The scRNA-seq data were obtained from public databanks (GSE84133, GSE85241, E-MTAB-5061, GSE83139, GSE81608) of human pancreatic single-cell transcriptomes. We first randomized this available pancreatic scRNA-seq transcriptomic data and allocated 80% of samples to a discovery/training set (Training; N=11,652 samples) and remaining into a validation/testing set (Test; N=2,913 samples) as outlined in **Figure 1**. With the availability of several ML algorithms (**Supplementary Table 1**), we probed the discovery dataset using 10 different ML workflows (**Figure 2A**) to identify features highly associated with the presence of insulin transcripts in a single cell. Genes (features) identified as the most important/predictive variables for each of these ML workflows were used to identify insulin transcript-containing cells from the validation set (remainder 20% of the samples). Validation results of the identified gene features from each of the 10 ML workflows are presented in the form of receiver operator characteristic (ROC) curves (**Figure 2B**). The top three ML algorithms; Gradient boosting, Random Forest and ADA boost (all Ensemble workflows), demonstrated similar performance returning an Area Under Curve (AUC) of between 0.83 – 0.86. A confusion matrix is presented below each ROC curve dataset (**Figure 2B**) to demonstrate the false-positive and false-negative predictions within every workflow. These analyses show that although Ensemble machine learning workflows are the best in predicting insulin-transcribing cells, other workflows, such as logistic regression, also perform closely to the Ensemble methods.

**Figure 1:**
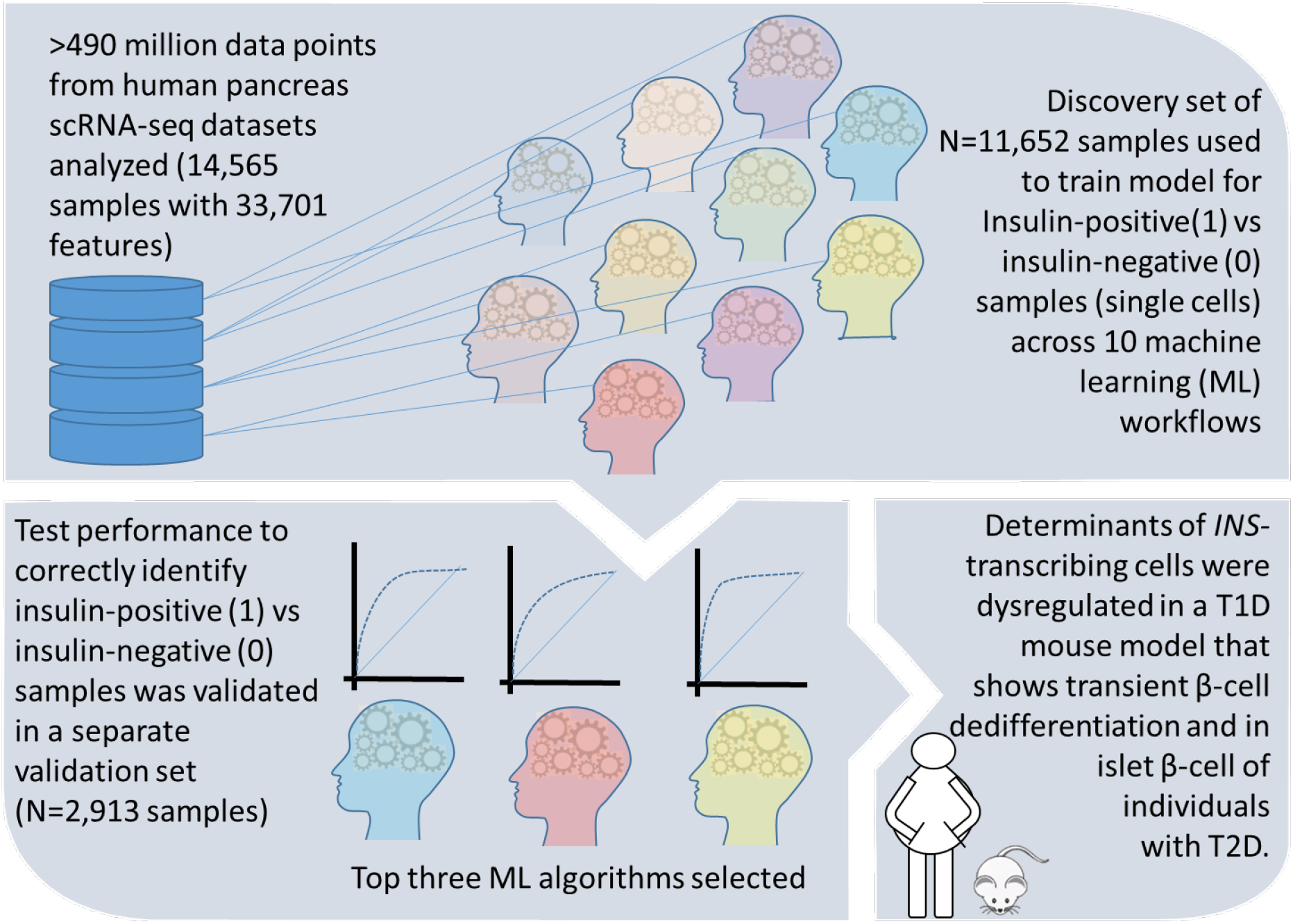
Comparative analysis of scRNA-seq datasets. Five different human pancreatic single-cell(sc)RNA-sequencing datasets (GSE84133, GSE85241, E-MTAB-5061, GSE83139, GSE81608) were curated to test the performance of 10 different machine learning (ML) algorithms. A training set comprising randomly selected 80% datasets (N=11,652 samples) were used in the learning phase (Training set; top panel). Ten ML workflows used in this study are symbolically illustrated as coloured heads. Each of the workflows aimed to identify a list of weighted features that accurately identify the presence (1; “insulin-positive”) or absence (0; insulin-negative) of insulin gene transcripts. Learning outcomes were validated (bottom-left panel) in 20% (N=2,913) of the total samples, and the top three ML algorithms providing the best area under the curve (AUC) in a receiver operator characteristic (ROC) curve analysis were identified. Top features (genes) associated with insulin gene transcription were derived from each ML-workflow and validated in other datasets. We analyzed the expression of these genes in the Ireα^β-/-^ mouse pancreatic scRNA-seq dataset (GSE144471) profiling dedifferentiating pancreatic β-cells as well as in human T2D islet scRNA-seq dataset (GSE154126).

**Figure 2:**
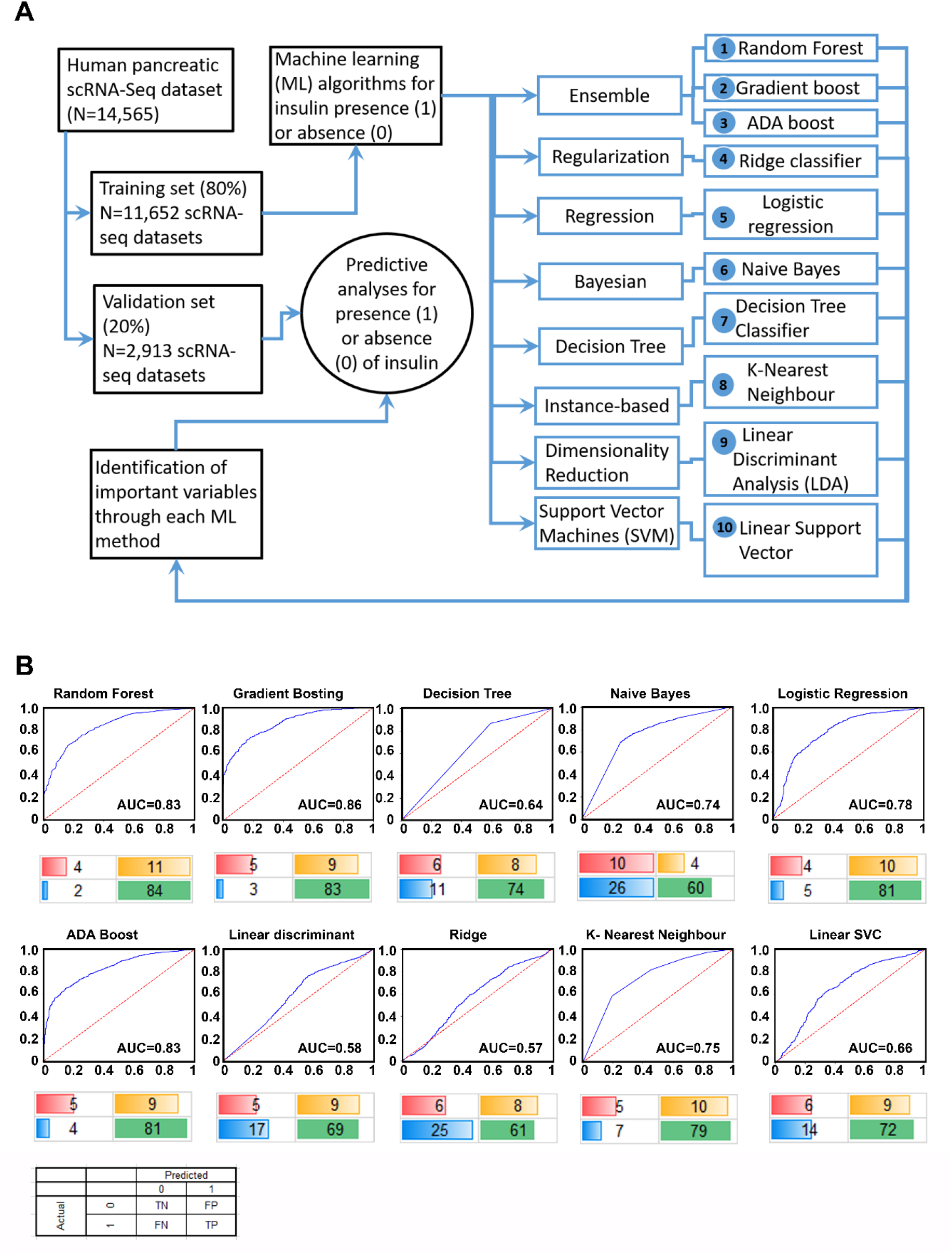
Study design and performance of different ML workflows. A flowchart of our analytical plan is presented in (**A**). Previously published datasets of single-cell RNA-sequencing analyses from pancreatic islet cell preparations were randomly divided into a training (N=11,652) and a validation (N=2,913) set. The learning phase (Training) involved identifying features (genes) and their associated weights/coefficients in each of the 10 machine learning (ML) methods (listed 1-10). Weighted features were used in the prediction of insulin transcription (across 10 ML algorithms) to test the performance of these models in an independent validation set of samples (N=2,913). ROC curve plots for each ML algorithm using validation set data are presented in (**B**). The area under the curve (AUC) for the tested workflows are presented along with a confusion matrix below the plot. Percent values are rounded off to the nearest integer (and hence may not sum up to an absolute 100%) and represent true negative (red), true positive (green), false positive (yellow) and false negative (blue) samples identified in the validation set.

### Ensemble ML workflows to identify genes associated with insulin transcription

The scRNA-seq datasets obtained from public databanks of human pancreatic single-cell transcriptomes were classified as insulin-transcribing (1) or those with no insulin (0) (**Figure 3A**). As described earlier, all the three Ensemble ML workflows presented with an AUC that was better than any of the other ML workflows tested in our ROC curve analysis. Ensemble workflows also presented with high accuracy (≥87%), precision (≥0.89), and sensitivity (≥0.95), which was comparable to other popular workflows such as logistic regression (**Figure 3B**). As Ensemble ML workflows such as Random Forest use a collection of decision trees (forest), we decided to compare the performance of the top three (Ensemble) workflows to a single (Decision tree) algorithm. The relative contribution of the top 10 features (genes) from each of these ML workflows are presented as radar plots (**Figure 3C**), whilst the longer list of genes ranked by their importance is presented in **Supplementary Table 2**. *IAPP, ADCYAP1, LDHA* and *SST* were common to all three Ensemble workflows. We then examined the pathways targeted by these features (genes) identified through each of the Ensemble and Decision Tree classifier by comparing them to a separate islet β-cell dataset (**Figure 3D**). Number of GO terms enriched across all four ML workflows (**Figure 3D**) suggests several common pathways (including insulin secretion) targeted by the features identified through these analysis (**Figure 3D, Supplementary Table 3**). These genes were also validated in a bulk RNA-seq dataset (GSE152111, n=66) of human islet samples (**Supplementary Figure 1**). In this analysis (**Supplementary Figure 1)**, most of these gene transcripts had significant positive correlation with insulin transcript. While some of the gene transcripts such as *LDHA, CRP, RPS15* and *RPL35* negatively correlated with insulin transcript in human islets.

**Figure 3:**
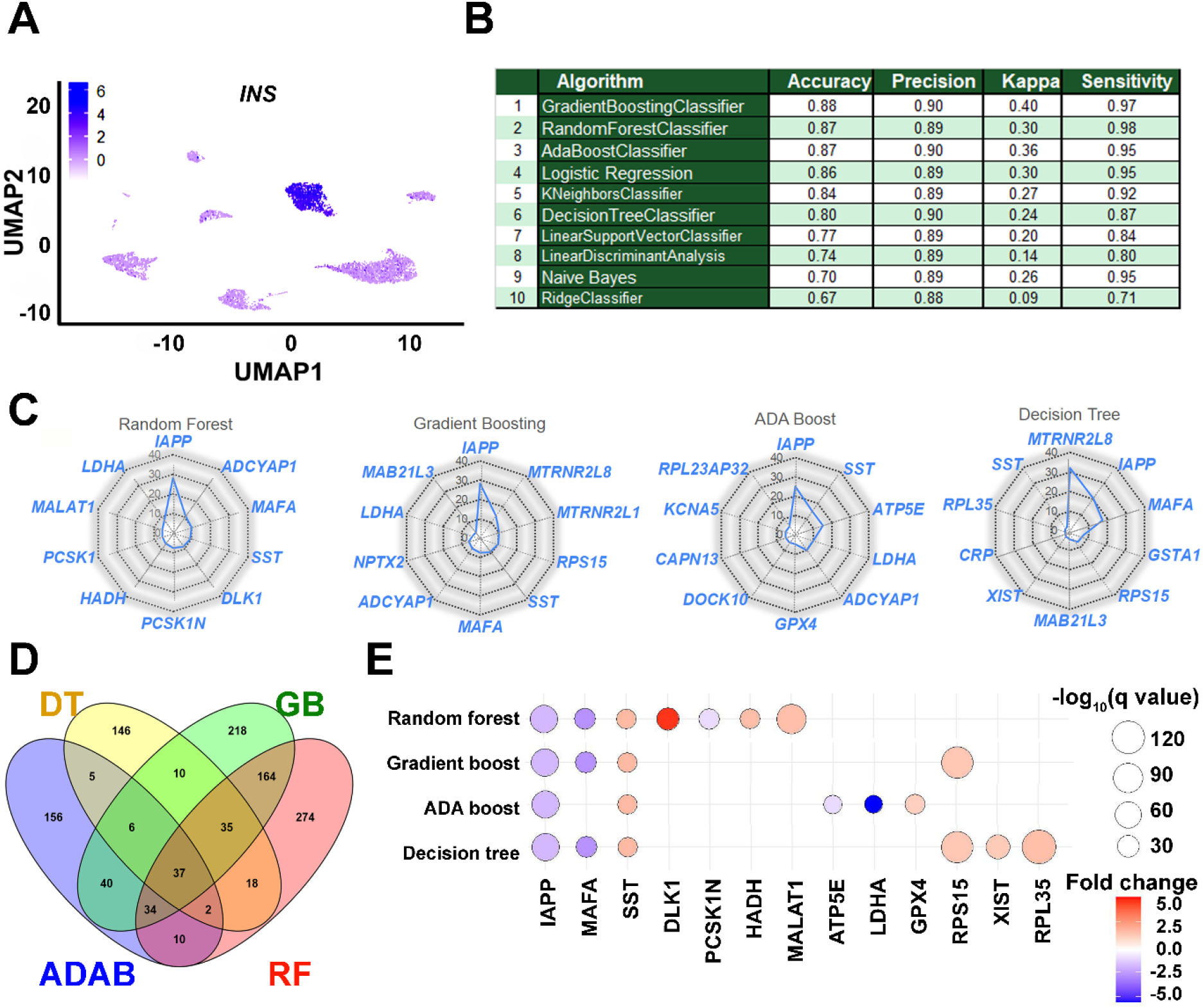
Performance and application of learned features in understanding insulin gene transcription. (**A**) A 2D clustering of pancreatic single cells assessed in this study using UMAP (Uniform Manifold Approximation and Projection plot). Cellular subtypes based on the UMAP clustering algorithm are labelled and graded (scale, inset) as per the level of insulin gene transcripts. (**B**) The performance of learning models on accurately identifying insulin-positive (1) and insulin-negative (0) single cells from the validation dataset are presented. (**C**) Relative weighted rank contributions of the top 10 genes in each of the four listed ML algorithms are presented as spider plots plotted in the order of importance (starting clockwise at 12-O’clock position). Percent representation of each of the genes indicates their relative contribution in the set on the spider plot with a logarithmic scale (center=1% and outer circle=100%). A comparison of the gene features identified by the top three ensemble workflows is presented along with those identified by the Decision Tree classifier. (**D**) Pathways targeted by up to the top 100 features (**Supplementary Table 2**) from each of the four selected ML methods (RF: Random Forest; GB: Gradient Boosting; ADAB: ADA Boost; DT: Decision Tree) identified using gene ontology (GO) function analysis are presented in the Venn diagram. Number of GO terms enriched and common for top features (genes) in each ML method are plotted. (**E**) All significantly dysregulated genes identified from and common to the four ML algorithms (**Figure 3C)** presented herein were assessed in the scRNA-seq dataset from Ire1α^β-/-^ mice. Bubble plot presenting fold-change and statistical significance (q-value) for each of the genes in Ire1α^fl/fl^ and Ire1α^β–/–^ mice are shown. Blue colour represents downregulation while red colour indicates increased abundance of transcripts in Ire1α^β-/-^ mice compared to control.

### Insulin-associated genes are dysregulated during β-cell dedifferentiation

Dedifferentiation of β-cells, characterized by the loss of expression of key β-cell maturation marker genes with an accompanying reduction in insulin secretion, has been observed in mouse models of type 1 (T1D) and type 2 (T2D) diabetes, as well as in individuals with diabetes[16-19]. We questioned if the expression of gene variables identified and validated (in silico) as being predictive of insulin gene transcription (**Figure 3C**) are dysregulated in a mouse model of T1D with evidence of islet dedifferentiation. Transient dedifferentiation of islet β-cells was recently reported in an established T1D preclinical mouse model upon β-cell-specific deletion of a key stress response gene, *Ire1α*, (Ire1α^β-/-^)[20]. These mice also demonstrated reduced β-cell number as well as diminished expression of insulin transcripts in β-cells compared to control (Ire-1α^fl/fl^) mice. Therefore, we evaluated the expression of the total 25 gene transcripts that made up the top 10 features across the four different ML workflows (**Figure 3C**) in the single cell datasets generated from these (Ire1α^β-/-^ and Ire1α^fl/fl^) islets. Twelve of these features were not significantly different between Ire1α^β-/-^ and Ire1α^fl/fl^ islets. However, the remaining thirteen features were significantly dysregulated in β-cells of Ire-1α^β-/-^ mice that were undergoing dedifferentiation (**Figure 3E**). Dedifferentiating β-cells showed significant downregulation of five key genes; *Iapp, MafA, Pcsk1n, Atp5e* and *Ldha*, whilst all other insulin-associated gene transcripts showed significantly higher levels (**Figure 3E**).

In type 2 diabetes (T2D), it is known that *INS* transcript expression is reduced. Therefore we validated the top, common gene features (*IAPP, SST, MAFA, ADCYAP1* and *LDHA*) from the three ML workflows using a separate publicly available single-cell RNA-seq dataset from non-diabetic (ND) vs T2D adult human pancreas (GSE154126[21]). Four of the five genes (*IAPP, SST, MAFA, ADCYAP1*), were significantly lower in T2D insulin-transcribing cells compared to ND insulin-transcribing cells (**Supplementary Table 4**).

## Discussion

In this study, we compared the performance characteristics of 10 different ML algorithms, (**Supplementary Table 1**) that are currently used in big data analyses. We analyzed a scRNA-seq dataset that was randomly split to a larger (80%; 392,684,052 data points) training set involving model learning, and then a smaller (20%; 98,171,013 data points) validation set. All algorithms identified a set of genes (features) that associate with insulin-production (1) defined as the presence of one or more transcripts of insulin in a sample, or no insulin production (0) from the 11,652 single cells analysed in the training test. We validated the predictive features identified through each ML workflow in the validation/test set of 2,913 single cell transcriptomes. ML workflows that returned high performance (based on AUC, sensitivity/specificity) were selected and the top 10 genes (ranked by their importance) in each of those ML methods were re-validated in discrete mouse and human datasets that model beta cell dedifferentiation (**Figure 3E, Supplementary Table 4**).

Our analysis provides two major outcomes that are of interest to a broad range of data analysts and biologists. First, a comparison of the ML algorithms identified Ensemble-based ML methods as the best performing algorithms in our analyses. Logistic regression performed closest to Ensemble methods, in line with previous reports in clinical datasets[22]. We then compared Ensemble methodologies to Decision tree algorithm. Decision tree offers the often-desired simplistic model generation method as compared to Ensemble methods such as Random Forest. The latter builds multiple decision trees independently and offers an overall learning model that is closest to the best possible prediction. Indeed, Decision tree was determined to be a weaker predictor than the Random Forest as the latter reduces variance using different sample sets (bootstrap) in training, randomizing feature subsets, and combining the predictive learning by building multiple decision trees. Random Forest prediction outcomes were similar to gradient boosting, which also builds a set of decision trees, but one tree at a time. The bagging and boosting approach used in ADA/Gradient boosting methods seems to have offered better accuracy and performance in insulin prediction analysis than those observed using Random Forest, whereas the Random Forest algorithm offered the highest sensitivity (**Figure 3B**) amongst all methodologies tested.

The other outcome from this analysis is the identification of genes that are associated with and predictive of insulin gene transcription in single cells. Since bulk RNA-sequencing studies do not offer the desired single-cell resolution to identify transcriptional regulation at a cellular level, our analyses provide a firsthand view of insulin gene transcriptional determinants identified through an unbiased, big data machine learning approach. The top three methodologies (based on high AUC values) belonged to Ensemble machine learning workflow. Weighted relative importance of the top-10 most important features are compared (**Figure 3C**). Interestingly, five genes were common to the top 10 features from all the algorithms compared – *IAPP, ADCYAP1, MAFA, SST* and *LDHA*. The top-ranked gene associated with insulin gene transcription across all the Ensemble workflows was *IAPP*. Islet amyloid polypeptide (*IAPP*) and insulin are known to be expressed in pancreatic islet β-cells and co-secreted in response to changes in glucose concentration[23, 24]. Their mRNA levels are also regulated by glucose. The promoters of both these genes share similar cis-acting sequence elements, and both bind the master regulatory transcription factor *PDX1*[23]. *FoxA2* (*HNF-3β*) negatively regulates *IAPP* promoter activity[25] and has also been shown to suppress insulin gene expression[26]. Although insulin gene expression is known to be regulated by several islet-enriched transcription factors, *MafA* is the most well recognized β-cell-specific activator of insulin gene expression[27]. The selection of *MAFA* as a key feature by three of the compared ML approaches tested through this analysis is therefore not surprising. The inclusion of *SST* in the top three gene features is intriguing. Somatostatin expression is known to be important in control of insulin release and ablation of somatostatin-expressing delta cells impairs pancreatic islet function and cause neonatal death in rodents[28]. SST analogs were shown to inhibit the release of insulin via the activation of both ATP sensitive K+ channels and G protein-coupled inward rectifier K+ channels[29]. Another candidate that was identified through these analyses is *MTRNR2L8*, a neuroprotective and antiapoptotic peptide derived from a portion of the mitochondrial *MT-RNR2* gene and reported in fetal as well as adult beta cells[30]. *ADCYAP1* stimulates insulin secretion in a glucose-dependent manner[31] and genetic screening in T2D Caucasians indicated the presence of two SNPs in exons 3 and 5 of this gene to be associated with T2D[32]. Finally, *LDHA*, which was also selected through these unbiased analyses across the top-three ML workflows is a pancreatic β-cell disallowed gene[33-35] and human *LDHA* levels are predictive of insulin transcription[36]. Consistent with these previous reports, our validation analysis in human islets RNA-seq data, demonstrated negative correlation of *LDHA* and positive correlation of *ADCYAP1, MAFA* and *SST* transcripts with insulin (**Supplementary Figure 1**). Together, these algorithms help in identifying a set of genes expressed in or disallowed from insulin-producing pancreatic β-cells.

Mouse models often provide the validation to understand mechanisms that cannot be tested in human studies. The Ire1α^β-/-^ mouse offers a unique model, wherein pancreatic β-cells transiently dedifferentiate during early post-natal life, allowing these knockout mice to escape immune-mediated β-cell destruction and T1D in later life[20]. Analysis of islet single cell sequencing data from this model identified genes that were significantly dysregulated in β-cells of Ire1αβ-/- mice when compared to control (Ire1α^fl/fl^) mice. Eight of thirteen features (from the top 10 features in each of the four ML workflows, **Figure 3C**), which showed significant dysregulation between Ire1α^β-/-^ and Ire1α^βfl/fl^ mice are upregulated in β-cells of Ire1α^β-/-^ mice (**Figure 3E**). Analysis of T2D islet single cell data also revealed down-regulation of four common gene features (*IAPP, SST, MAFA* and *ADCYAP1* identified across our three top ML workflows) in T2D compared to ND insulin transcribing cells (**Supplementary Table 4**). Interestingly, Delta Like Non-Canonical Notch Ligand 1 (*DLK1*) was also significantly downregulated in T2D compared to ND insulin transcribing cells (Mann-Whitney test P-value=0.0017). The imprinted region of chromosome 14q32.2, contains microRNA cluster of *DLK1*-*MEG3* which are highly expressed and more specific in human β-cells compared to α-cells. Previous study had also shown that in T2D human islets, the *MEG3*-microRNA locus expression levels are significantly lower[37]. The 14q32 locus of microRNAs (such as co-expression of miR-376a and miR-432) also have been shown to target and suppress the expression of *IAPP*[37], that was one of the top features in our analyses.

### Strength and Limitation

This is a first demonstration comparing multiple ML algorithms to identify key genes associated with insulin transcription using a big dataset of over 490 million data points. As anticipated, Ensemble methods perform better than most other workflows and identified a set of genes that corroborate with previous reports of transcriptional regulation of insulin in mouse and human β-cells. These findings indicate that unbiased ML workflows for big data analyses can generate biologically meaningful results, when applied to large training datasets. Our study provides the codes/scripts for other researchers to use in existing as well as emerging datasets for identification of gene candidates associated with other genetic pathways (eg. related to *GCG* or *GCK*) in future or to genes recognized to be associated with T2D GWAS datasets. We recognize that there are several limitations: we are unsure as to why some other well known candidates (such as *PDX1*, and *NEUROD*) were not selected by our top predictive models. An explanation is that we used a whole pancreatic single cell dataset and that the predictive models generated through filtering out β-cells may be more enriched for known pro-endocrine gene regulators such as *PDX1*. The other explanation is that although *PDX1* is a key regulator, the transcript levels in these datasets using multiple scRNA-seq technologies may not be sufficient considering the sequencing depth offered by some of these scRNA-seq workflows.

We recognize that exhaustive (eg. LOOCV,[38]) as well as non-exhaustive cross-validation approaches (such as K-fold cross-validation [39]) were not performed here. Such cross-validation approaches, although useful in assessing how results will generalize to an independent dataset, are mostly used in the validation of much smaller datasets. In big data analyses, the use of such cross-validation methodologies would limit the analyses to only those with an access to high-end cluster computing. The 10 different ML scripts used in these analyses are designed to work on a high-end personal computing device (i7 processor with 4 cores and 32GB RAM or better). We believe that the application of such ML algorithms to the expanding scRNA-seq datasets would lead to the confirmation/validation of current as well as identification of determinants of gene transcription, thereby accelerating innovation in discovery of gene targets in biology and medicine.

## Methods

### Pancreatic single-cell (sc)RNA sequencing datasets and analyses

#### Human pancreatic single-cell sequencing datasets

The pancreatic single-cell sequencing dataset (N=14,890) was extracted using the Panc8 data containing multiple publicly available scRNA-seq transcriptomes (GSE84133, GSE85241, E-MTAB-5061, GSE83139, GSE81608). Analysis was carried out by using R studio version 1.2.5033, the SeuratData (version 0.2.1) and Seurat (3.2.3). Data normalization across multiple datasets in Panc8 are described previously[40]. Single-cell selection criteria involved all cells that contained transcript data (samples with all zeros eliminated). Single-cell datasets were randomly split to 80% samples selected in a training set with the remaining 20% retained for validation. ML-based identification of features predicting insulin transcript was carried out by data scientists blinded to gene variables and sample information. Analytical plans (80/20 discovery and validation) were predetermined and gene identifiers/sample details were provided once models were ranked following in silico validation.

#### Ire1α^β-/-^ mouse pancreatic single-cell dataset

Single-cell RNA-seq dataset from pancreatic islets of Ire1α^fl/fl^ (N=1,163 single-cell transcriptomes from one mouse) and Ire1α^β–/–^ (N=1,683 single-cell transcriptomes from two mice) were obtained through GSE144471[20]. The β-cells (Ire1α^fl/fl^: 830 cells; Ire1α^β–/–^: 816 cells) were separated from the dataset and the expression values of selected genes were evaluated in the β-cell population.

#### T2D pancreatic single-cell (sc)RNA sequencing dataset

Pancreatic single cell normalized read dataset of adult ND (with no diabetes; N=4) and T2D (N=10) donors were obtained from GSE154126 [21]. The adult ND (N=296) and T2D (N=505) insulin transcribing cells were compared and used for validation.

#### Human pancreatic single-cell sequencing classification and analyses

Deidentified datasets were shared with data scientists. A random number generator function was used to allocate 80% of samples to a training set. Analyses were carried using Python (Ver:3.4), wherein the data file was imported into the data frame, transposed, edited to delete INS and INS-IGF2 columns from the data frame and labelled (label=0 where INS=0 and label=1 where INS>0). Classifiers were initialized and model trained using the discovery (80%) data set. Predictive analyses were then carried out on the validation (20%) set and the resulting accuracy metrics were saved to compare the feature importance. Selected classifiers (Random Forest, Gradient Boosting, Decision Tree Classifier, Logistic Regression, Multinomial Naive Bayes Classifier, ADA Boost Classifier, Linear Discriminant Analysis, Ridge Classifier, KNeighbors Classifier and Linear Support Vector Classifier) were used on the same set and codes for these analyses will be made available through GitHub on publication.

### Pancreatic islet RNA sequencing dataset and analysis

#### Human pancreatic islet bulk RNA sequencing dataset

Human pancreatic islet RNA-seq dataset was obtained from GSE152111 [41]. RNA-seq dataset contains n=66 human islet samples, across 65 different donors with no diabetes. Two of the 66 RNA-seq samples were duplicates from the same donor and their RNA-seq profile highly correlative (Pearson r=0.99) to each other. The average of the duplicates of this donor was calculated prior to analysis. Data was analyzed in DEseq values.

### Pathway analysis

To analyze enrichment for β-cell pathways, lists of pancreatic single-cell features generated/predicted by ML algorithms (Random forest, Gradient boosting, Decision tree classifier and ADA Boost classifier) were compared with β cell-expressed genes (E-GEOD-20966) using Gene Ontology (GO) over-representation analysis on Pantherdb.org[42]. Preanalytical workflows included cleaning up entries not mapping to protein-coding gene symbols in E-GEOD-20966. Gene lists for each ML algorithm consisted of up to the top 100 genes as predictors of insulin expression, which were compared against the data set of β-cell expressed genes (N=13,165 from E-GEOD-20966). Overrepresentation analysis using GO categories for biological processes (GO: BP) was performed using binomial testing using false-detection-rate to correct for multiple testing. Lists of significantly enriched pathways associated with each ML algorithm were compared using Venn diagrams[43].

### Statistical analysis

The R software (ver. 3.6.1; R Foundation for Statistical Computing, Vienna, Austria) was used to create the categorical bubble plot using the packages ggplot2 (3.3.3), ggpubr (0.4.0) and proto (1.0.0). Spearman correlation matrix analysis was generated through using R packages corrplot (0.90), Hmisc (4.6.0), dplyr (1.0.7) and readxl (1.3.1) in R and Rstudio software. Statistical software, Microsoft Excel (ver. 2016; Microsoft, Redmond, WA, USA), the R software and/or GraphPad Prism (ver. 8.4.1; GraphPad Software, San Diego, CA, USA) were used for univariate test comparisons and Benjamini-Hochberg method for multiple testing.

## Resource Availability

Lead Contact: AAH.

## Code Availability

The data codes/scripts will be made available through this publication to carry similar analyses in these and related datasets. The codes will be made available through https://github.com/Isletbiology/ML.

## Author contributions

Conceptualization: AAH; Methodology: AAH, AP, SB, LTD; Software: VT, MVJ, WKMW, CXD, HL, FE, LTD; Validation: WKMW, MVJ, VT, CXD, AB; Data Curation: WKMW, MVJ, VT, CXD, HL, YVC, WJH, FE, LTD, AAH; Writing – Original Draft: AAH, MVJ, WKMW, LTD; Review & Editing: All authors; Visualization: AAH; Supervision: AAH; Project Administration: AAH; Funding Acquisition: AAH.

## Acknowledgements

AAH acknowledges the initial interactions with Tune Pers, University of Copenhagen, Denmark.

## Supplementary information

**Supplementary Table 1:** Description of the ten different ML workflows used for predictive analyses. Related to Figure 2A.

| # | Workflow | Build | Description |
| --- | --- | --- | --- |
| 1 | **Random Forest** | Ensemble | Random Forest classifier is a meta-estimator that fits several decision trees on various sub-samples of datasets and uses an average to improve the predictive accuracy of the model and controls over-fitting. The sub-sample size is always the same as the original input sample size, but the samples are drawn with replacement. Reduction in over-fitting and Random Forest classifier is more accurate than decision trees in most cases. |
| 2 | **Gradient Boosting** | Ensemble | The idea behind "gradient boosting" is to take a weak hypothesis or weak learning algorithm and make a series of tweaks to it that will improve the strength of the hypothesis/learner. This type of Hypothesis Boosting is based on the idea of Probability Approximately Correct Learning (PAC). |
| 3 | **Adaptive Boosting (Adaboost)** | Ensemble | For AdaBoost, many weak learners are created by initializing many decision tree algorithms that only have a single split. The instances/observations in the training set are weighted by the algorithm, and more weight is assigned to instances that are difficult to classify. More weak learners are added into the system sequentially, and they are assigned to the most difficult training instances. In AdaBoost, the predictions are made through majority vote, with the instances being classified according to which class receives the most votes from the weak learners. |
| 4 | **Ridge Classifier** | Regularization | Ridge classifier first converts binary targets to [-1, 1] and then treats the problem as a regression task, optimizing the same objective as above. The predicted class corresponds to the sign of the regressor's prediction. For multiclass classification, the problem is treated as multi-output regression, and the predicted class corresponds to the output with the highest value. |
| 5 | **Logistic Regression** | Regression | Logistic regression is a machine learning algorithm for classification. In this algorithm, the probabilities describing the possible outcomes of a single trial are modelled using a logistic function. It is most useful for understanding the influence of several independent variables on a single outcome variable. |
| 6 | **Naive Bayes** | Bayesian | Naive Bayes algorithm based on Bayes' theorem with the assumption of independence between every pair of features. Naive Bayes classifiers work well in many real-world situations such as document classification and spam filtering. This algorithm requires a small amount of training data to estimate the necessary parameters. Naive Bayes classifiers are extremely fast compared to more sophisticated methods. |
| 7 | **Decision Tree Classifier** | Decision Tree | Given data of attributes together with its classes, a decision tree produces a sequence of rules that can be used to classify the data. Decision Tree is simple to understand and visualise, requires little data preparation, and can handle both numerical and categorical data. |
| 8 | **K-Nearest Neighbours** | Instance-based | Neighbours based classification is a type of lazy learning as it does not attempt to construct a general internal model but simply stores instances of the training data. Classification is computed from a simple majority vote of the k nearest neighbours of each point. This algorithm is simple to implement, robust to noisy training data, and effective if training data is large. |
| 9 | **Linear Discriminant Analysis (LDA)** | Dimensionality Reduction | It is a classifier with a linear decision boundary, generated by fitting class conditional densities to the data and using Bayes' rule. The model fits a Gaussian density to each class, assuming that all classes share the same covariance matrix. The fitted model can also be used to reduce the dimensionality of the input by projecting it to the most discriminative directions, using the transform method. |
| 10 | **Linear Support Vector Classifier** | Support Vector Machines (SVM) | Support vector machine is a representation of the training data as points in space separated into categories by a clear gap that is as wide as possible. New examples are then mapped into that same space and predicted to belong to a category based on which side of the gap they fall. |

**Supplementary Table 2:**
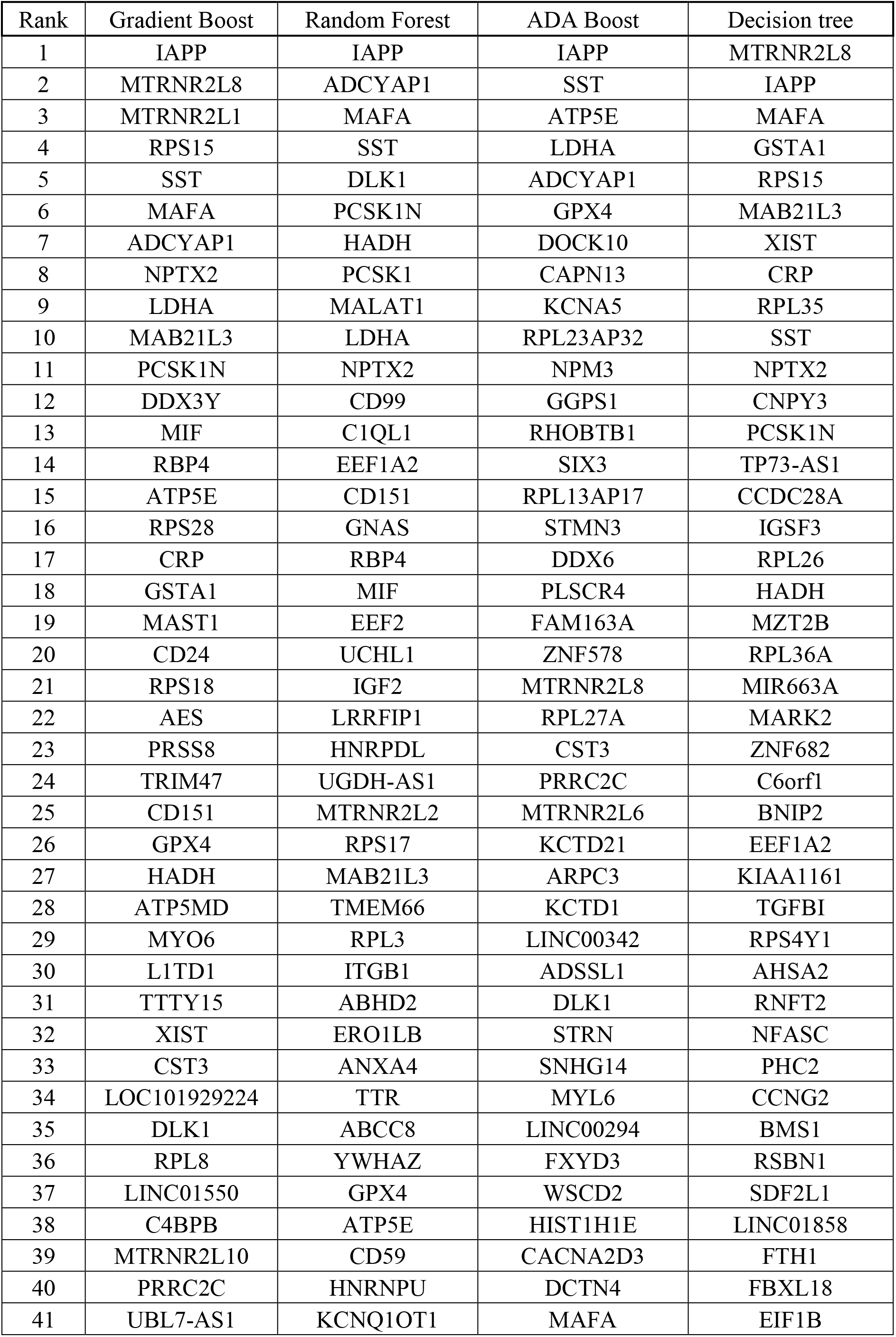
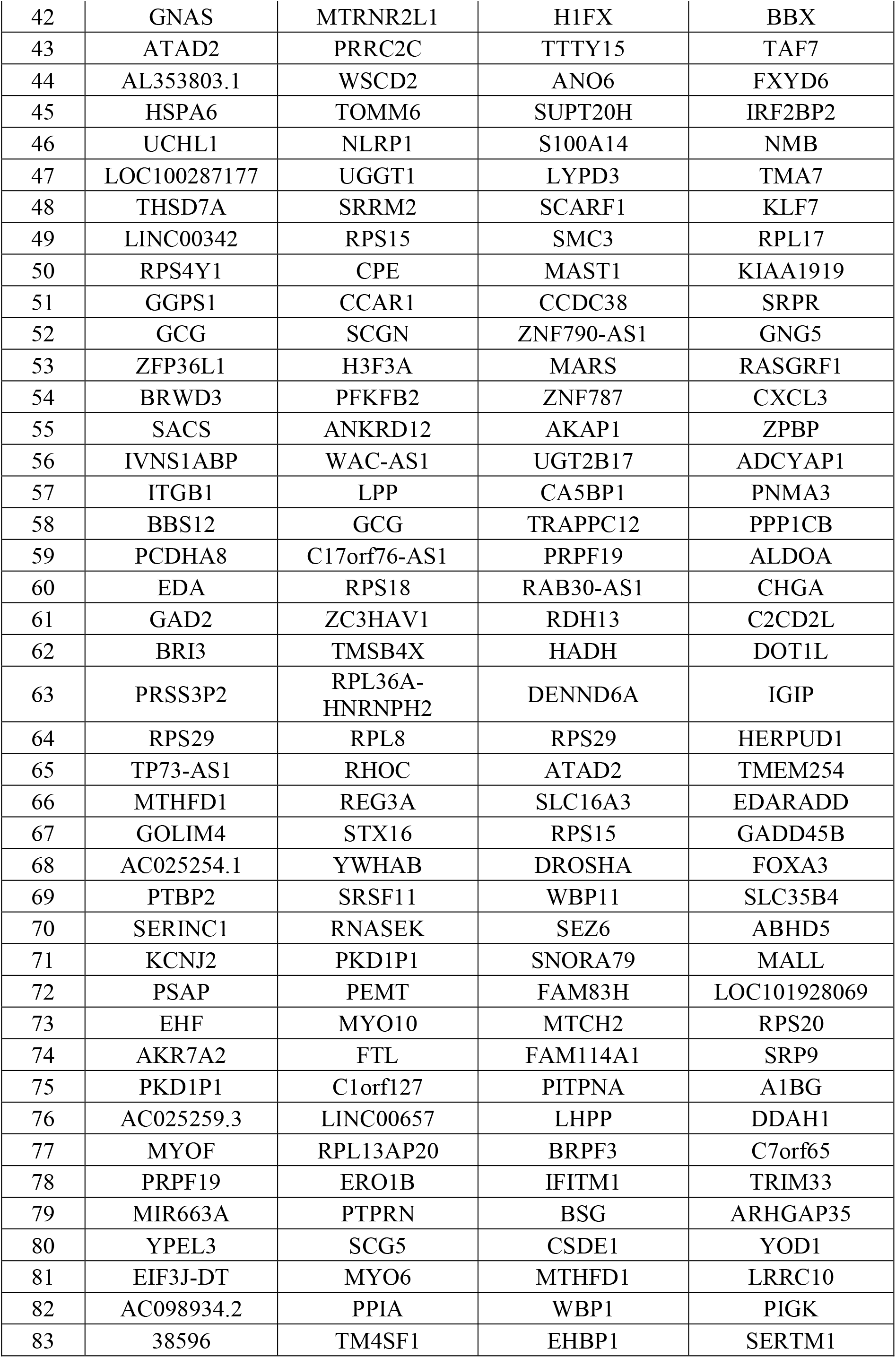
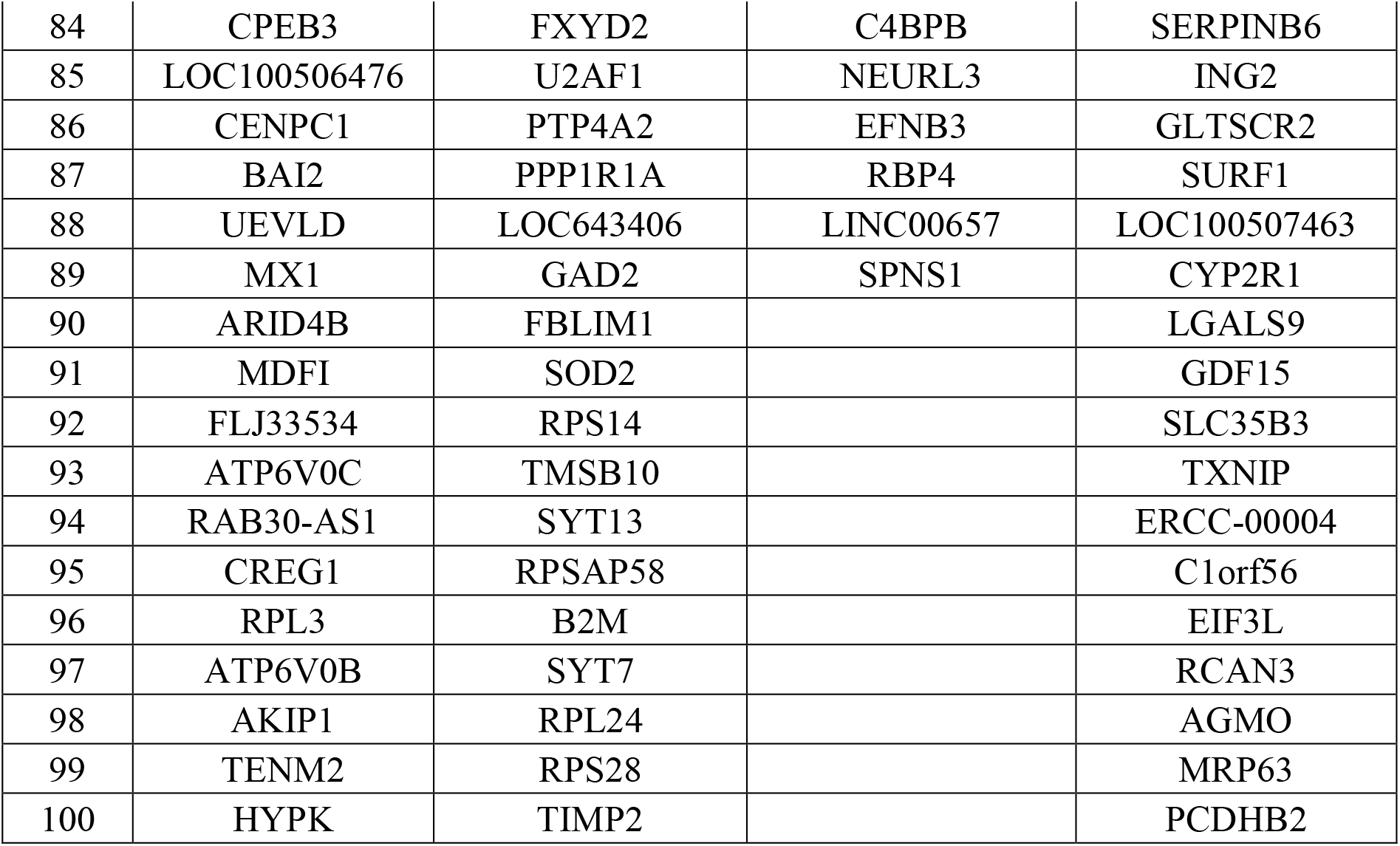
List of up to top 100 genes ranked by their importance in association with presence of insulin transcript in a single cell, identified from the four selected ML methods (Gradient Boosting; Random Forest; ADA Boost; Decision Tree). Related to Figure 3D.

**Supplementary Table 3:** The 37 common pathways (elements) targeted by the top 10 features from each of the four selected ML methods (RF: Random Forest; GB: Gradient Boosting; ADAB: ADA Boost; DT: Decision Tree) identified using gene ontology (GO) function analysis. Related to Figure 3D. List ordered in alphabetical order.

| # | **37 common elements in "AdaBoost", "DecTree", "GraBoost" and "RanForest":** |
| --- | --- |
| 1 | amylin receptor signaling pathway (GO:0097647) |
| 2 | calcitonin family receptor signaling pathway (GO:0097646) |
| 3 | cardiolipin acyl-chain remodeling (GO:0035965) |
| 4 | cotranslational protein targeting to membrane (GO:0006613) |
| 5 | establishment of protein localization to endoplasmic reticulum (GO:0072599) |
| 6 | fatty acid beta-oxidation (GO:0006635) |
| 7 | histamine secretion (GO:0001821) |
| 8 | histamine transport (GO:0051608) |
| 9 | hormone-mediated apoptotic signaling pathway (GO:0008628) |
| 10 | insulin secretion (GO:0030073) |
| 11 | mRNA catabolic process (GO:0006402) |
| 12 | negative regulation of acute inflammatory response (GO:0002674) |
| 13 | negative regulation of acute inflammatory response to antigenic stimulus (GO:0002865) |
| 14 | negative regulation of acute inflammatory response to non-antigenic stimulus (GO:0002878) |
| 15 | negative regulation of amyloid fibril formation (GO:1905907) |
| 16 | nuclear-transcribed mRNA catabolic process (GO:0000956) |
| 17 | nuclear-transcribed mRNA catabolic process, nonsense-mediated decay (GO:0000184) |
| 18 | peptide hormone secretion (GO:0030072) |
| 19 | positive regulation of calcium ion import across plasma membrane (GO:1905665) |
| 20 | positive regulation of somatostatin secretion (GO:0090274) |
| 21 | protein targeting to ER (GO:0045047) |
| 22 | regulation of acute inflammatory response to non-antigenic stimulus (GO:0002877) |
| 23 | regulation of amyloid fibril formation (GO:1905906) |
| 24 | regulation of calcium ion import across plasma membrane (GO:1905664) |
| 25 | regulation of hormone levels (GO:0010817) |
| 26 | regulation of hormone secretion (GO:0046883) |
| 27 | regulation of oligodendrocyte progenitor proliferation (GO:0070445) |
| 28 | regulation of peptide hormone secretion (GO:0090276) |
| 29 | regulation of somatostatin secretion (GO:0090273) |
| 30 | response to extracellular stimulus (GO:0009991) |
| 31 | RNA catabolic process (GO:0006401) |
| 32 | sensory perception of pain (GO:0019233) |
| 33 | spermine biosynthetic process (GO:0006597) |
| 34 | spermine metabolic process (GO:0008215) |
| 35 | SRP-dependent cotranslational protein targeting to membrane (GO:0006614) |
| 36 | viral gene expression (GO:0019080) |
| 37 | viral transcription (GO:0019083) |

**Supplementary Table 4:**
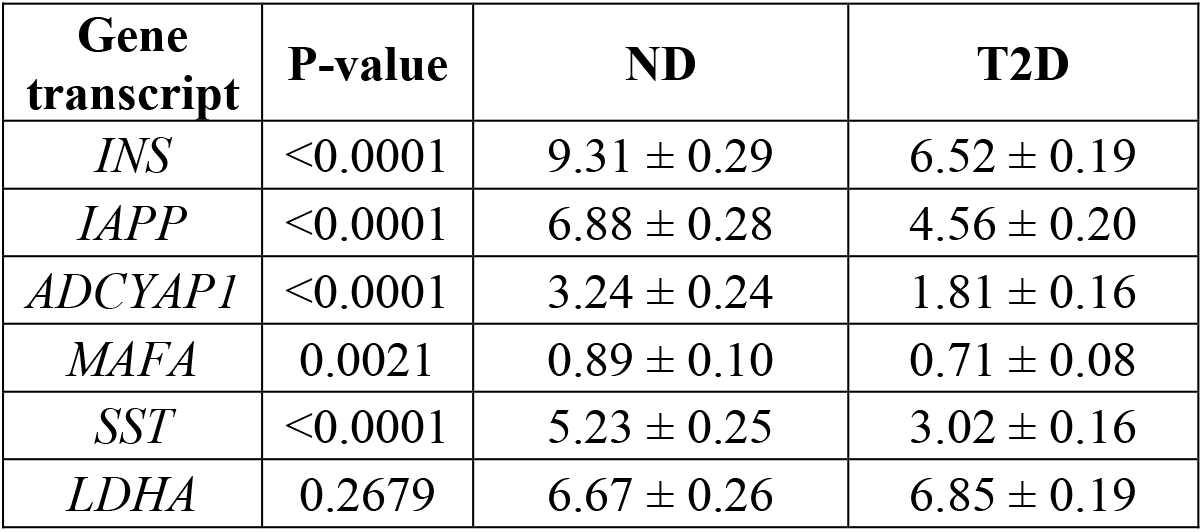
The key genes common in all top three ML workflows (Figure 3C) with *INS* gene accessed in single-cell RNA-seq data (GSE154126[21]) between adult non-diabetic (ND; N=296) vs T2D (N=505) insulin transcribing single cells. Data are presented as log_2_ transformed normalized reads mean ± SEM. The P-values were calculated with the Mann-Whitney test. Genes with significant P-value (P≤0.05) show here, remain significant after adjusting for multiple testing using the Benjamini-Hochberg method (FDR=0.05, on 21,153 tests).

**Supplementary Figure 1:**
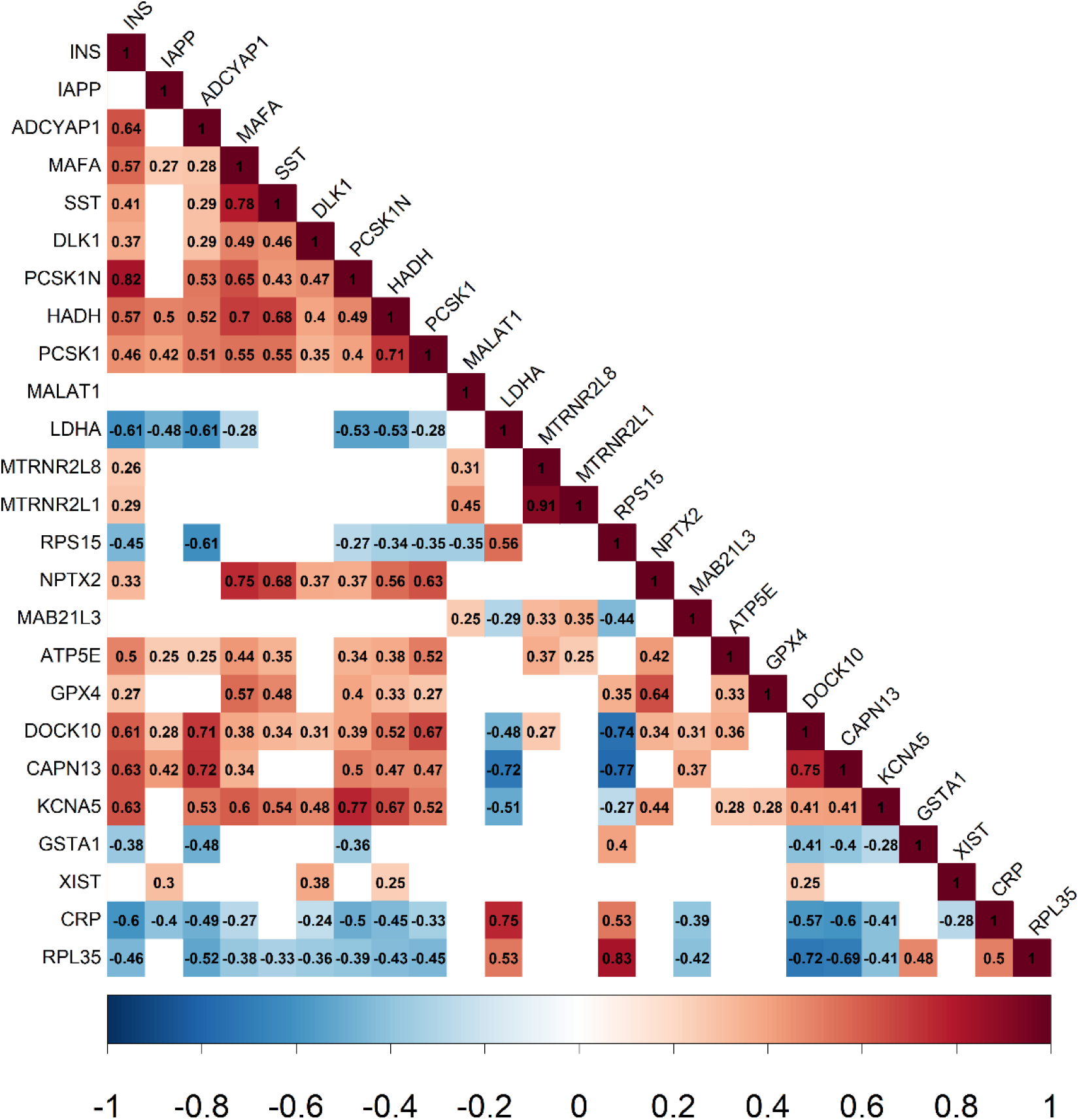
Selected genes validated in human islet bulk-RNA-seq dataset. Spearman correlation matrix of the top gene transcripts selected in the four ML algorithms (as presented in Figure 3C) along with insulin transcript on the human islet bulk RNA-seq dataset (GSE152111; n=66). RPL23AP32 was not present in the dataset. Only significant correlation (Spearman p-value < 0.05) are presented with colour along with the Spearman correlation r coefficient in black text within the square. Positive correlation is shown in shades of red, while negative correlation is in shades of blue.

